# Plasmid–plasmid interactions reshape extracellular vesicle cargo and gene transfer potential

**DOI:** 10.64898/2026.04.28.721519

**Authors:** Faina Tskhay, Haining Huang, Robin Starke, Magali de la Cruz Barron, M. Pilar Garcillán-Barcia, Thomas U. Berendonk, Anja Worrich, Uli Klümper

**Author notes:** corresponding authors Corresponding authors Dr. Uli Klümper, TU Dresden, Institute of Hydrobiology, 01062 Dresden, Zellescher Weg 40, Germany, Dr. Anja Worrich, BTU Cottbus-Senftenberg, Institute of Biotechnology, 01968 Senftenberg, Universitätsplatz 1, Germany. These authors contributed equally (Faina Tskhay, Haining Huang).

## Abstract

Plasmids are key drivers of horizontal gene transfer, yet their dissemination is not limited to conjugation. Extracellular vesicles (EVs) can transport plasmid DNA, but the factors governing plasmid incorporation into EVs remain poorly understood. Here, we tested whether principles of conjugative plasmid transfer, including plasmid mobility type and plasmid–plasmid interactions, extend to EV-mediated export. Using a conjugative plasmid (pKJK5) and a mobilizable plasmid (RSF1010) in two Gram-negative hosts, we quantified plasmid incorporation into EVs under single- and dual-plasmid conditions. When present individually, the conjugative plasmid was preferentially incorporated into EVs, exceeding RSF1010 by 10-23-fold despite its lower intracellular abundance. Under co-residence, this pattern reversed: RSF1010 became enriched by 13-39-fold, while pKJK5 was reduced by 2-7-fold. Consequently, EV-associated plasmid cargo shifted to RSF1010 dominance, deviating strongly from the expected 10-fold higher pKJK5 cargo if a stochastic model based on intracellular abundance and single-plasmid conditions were applicable. We propose that mobilizable plasmids under coexistence exploit conjugative plasmid transfer machinery to access membrane-associated sites, increasing their likelihood of incorporation into EVs. Our findings demonstrate that plasmid-plasmid interactions reshape EV cargo and identify a previously unrecognized mechanism that may influence extracellular gene transfer potential in microbial communities.

## Introduction

Bacterial extracellular vesicles (EVs) are membrane-derived, spherical nanoparticles with a size range of 20-300 nm that are released from the cell envelope through processes such as membrane blebbing^1–3^. Once considered mostly involved in bacterial waste disposal, EVs are now recognized as active contributors to bacterial survival and adaptation^4–6^. The versatility of EVs is linked with their ability to encapsulate, transport, and release different cargo molecules to mediate intercellular communication, facilitate pathogenicity, deliver bioactive compounds, or protect bacterial cells against antibiotics^4,5,7,8^. Beyond the delivery of proteins and metabolites, EVs can also encapsulate and transfer genetic material, such as free nucleic acids and plasmids^5,8–11^. These properties position EVs as a potential fourth mechanism of horizontal gene transfer (HGT)^10,12–17^.

In bacteria, HGT is mediated by three “classical” mechanisms: conjugation, transformation, and transduction^15,18,19^. However, unlike these conventional HGT mechanisms, EV-mediated transfer does not require cell contact, as in conjugation, nor is it limited by host specificity, as in transduction or recipient competence, as in transformation^10,16,20^. In addition, vesicular encapsulation protects the genetic cargo from endonuclease degradation, thereby enabling long-distance and even inter-species transfer of genetic material^8,10^. Given the central role of HGT in the spread of antimicrobial resistance (AMR) and virulence determinants within bacterial communities, additional mechanisms of HGT require further investigation^21–23^. Successful encapsulation and transfer of both mobile and chromosomal genetic elements through bacterial EVs have already been reported in numerous studies^8,10,14,24^. For example, EV-mediated transfer of β-lactamase-encoding plasmids in *Acinetobacter baylyi* resulted in the acquisition of ampicillin resistance by the recipient bacteria^8^.

Within the context of EV-mediated HGT, selective incorporation of specific gene clusters and genetic elements into EVs has been reported, suggesting that EV cargo is not randomly assembled^14,24^. For example, selective packaging of plasmids, key vehicles of HGT in bacterial communities^25,26^, into EVs could majorly affect how efficiently plasmid-encoded genes, such as ARGs, disseminate. The efficiency of plasmid packaging into EVs has previously been linked to plasmid-specific properties such as size, origin of replication, and intracellular copy number. Comparative studies have shown that high-copy-number plasmids are more frequently detected in vesicles than low-copy-number plasmids, suggesting that intracellular abundance contributes to packaging rates^10,27^. However, these observations are largely based on bacterial strains harbouring a single plasmid.

In addition to intracellular abundance, the spatial organization of plasmids within the cell may influence their incorporation into EVs. Conjugative plasmids encode mobility (MOB) functions and membrane-associated mating pair formation (MPF) complexes required for their autonomous transfer between bacteria^13,28^. These systems include type IV secretion machinery and coupling proteins that recruit plasmid DNA to the membrane-associated transfer complex, positioning it in proximity to the cell envelope^29,30^. In contrast, mobilizable plasmids encode only the MOB functions but lack the MPF complex and therefore depend on exploiting conjugative plasmids’ MPF complexes for transfer^13,22,25^. As a result, mobilizable plasmids are not inherently associated with membrane-bound transfer systems. Given that EVs originate from the cell envelope^2,4,5,31^, these differences in membrane association may influence plasmid incorporation into EVs. However, existing observations are largely based on single plasmid systems and do not address how such relationships operate in more complex intracellular contexts.

In natural microbial communities, bacteria frequently carry multiple plasmids, and interactions between coexisting plasmids are common^13,22,25,32^. Such interactions are well described for conjugative transfer, where mobilizable plasmids exploit conjugative machinery and can efficiently compete for access to type IV coupling proteins and membrane-associated transfer complexes^28^. These dynamics demonstrate that plasmid transfer is not solely determined by plasmid abundance or intrinsic properties, but also by interactions that affect access to membrane-associated systems. Whether plasmid–plasmid interactions between conjugative and mobilizable plasmids extend beyond conjugation to influence EV-mediated gene export remains unknown. Addressing this question is essential for understanding how EV cargo composition is determined under conditions that more closely reflect natural plasmid co-residence.

Taking into account the association of conjugative plasmids with the membrane and the high occurrence of plasmid interactions, which may influence incorporation of plasmids into EVs, we formulated two hypotheses: First, conjugative plasmids may be preferentially incorporated into EVs compared to mobilizable plasmids under single-plasmid conditions. Second, plasmid co-residence alters plasmid incorporation into EVs, consistent with reshaping access to and competition for positioning at membrane-associated transfer systems.

To test these hypotheses, we investigated the incorporation of a model conjugative low-copy-number plasmid (pKJK5) and a model mobilizable high-copy-number plasmid (RSF1010) into bacterial EV cargo across single- and dual-plasmid carriage states. We find that plasmid type and co-residence jointly shape EV cargo composition. While conjugative plasmids are preferentially incorporated into EVs in isolation, this pattern is altered under co-residence, where the mobilizable plasmid becomes enriched, and the conjugative plasmid is relatively depleted. This shift is consistent across different bacterial hosts, indicating that plasmid-plasmid interactions reshape EV-mediated gene export and represent a key determinant of EV cargo composition as well as a previously underappreciated layer of control in HGT.

## Materials and Methods

### Bacterial strains

To test how plasmid type and plasmid co-residence influence incorporation into extracellular vesicles, we used two Gram-negative host backgrounds, *Escherichia coli* and *Pseudomonas putida*, and two model plasmids with distinct transfer strategies: the conjugative IncP-1 plasmid pKJK5^33^ with a size of 57,419 base pairs and the 11,720 base pair, mobilizable IncQ plasmid RSF1010^34^, both carrying a GFP cassette ^35,36^.

For each host, four strain types were required to address the hypotheses: a plasmid-free strain, strains carrying either pKJK5 or RSF1010 individually, and a strain carrying both plasmids. All strains were derived from *E. coli* K12::*lac*I^q^-pLpp-*mCherry*-Km^R^ and *P. putida* KT2440::*lac*I^q^-pLpp-*mCherry*-Km^R^ chromosomal backgrounds, as in these strains GFP expression is repressed and thus not a significant cost factor. The plasmid-free and single-plasmid strains were already available from previous studies^35,36^. Therefore, to complete the set, we constructed strains carrying both plasmids in each host background.

To generate *E. coli* and *P. putida* strains carrying both plasmids, liquid mating was performed between the *E. coli* MG1655 carrying the RSF1010 plasmid that confers resistance to streptomycin and *P. putida* KT2440 carrying the pKJK5 plasmid that confers tetracycline resistance^35^. Overnight cultures of these strains were adjusted to OD_600_ = 0.5, mixed in a 1:1 ratio, and incubated overnight in Luria-Bertani (LB) medium (Merck KGaA, Darmstadt, Germany) at 120 rpm and 30°C. Successful acquisition of both plasmids was confirmed by plating 100 μL of the resulting mixed culture on selective media: Chromocult coliform agar for *E. coli* and Pseudomonas isolation agar (both Merck KGaA) for *P. putida*, supplemented with both antibiotics, streptomycin (100 µg/mL) and tetracycline (10 µg/mL). Together, these strains provided the full set of plasmid configurations required to test the hypotheses across both host backgrounds (Table 1).

**Table 1:**
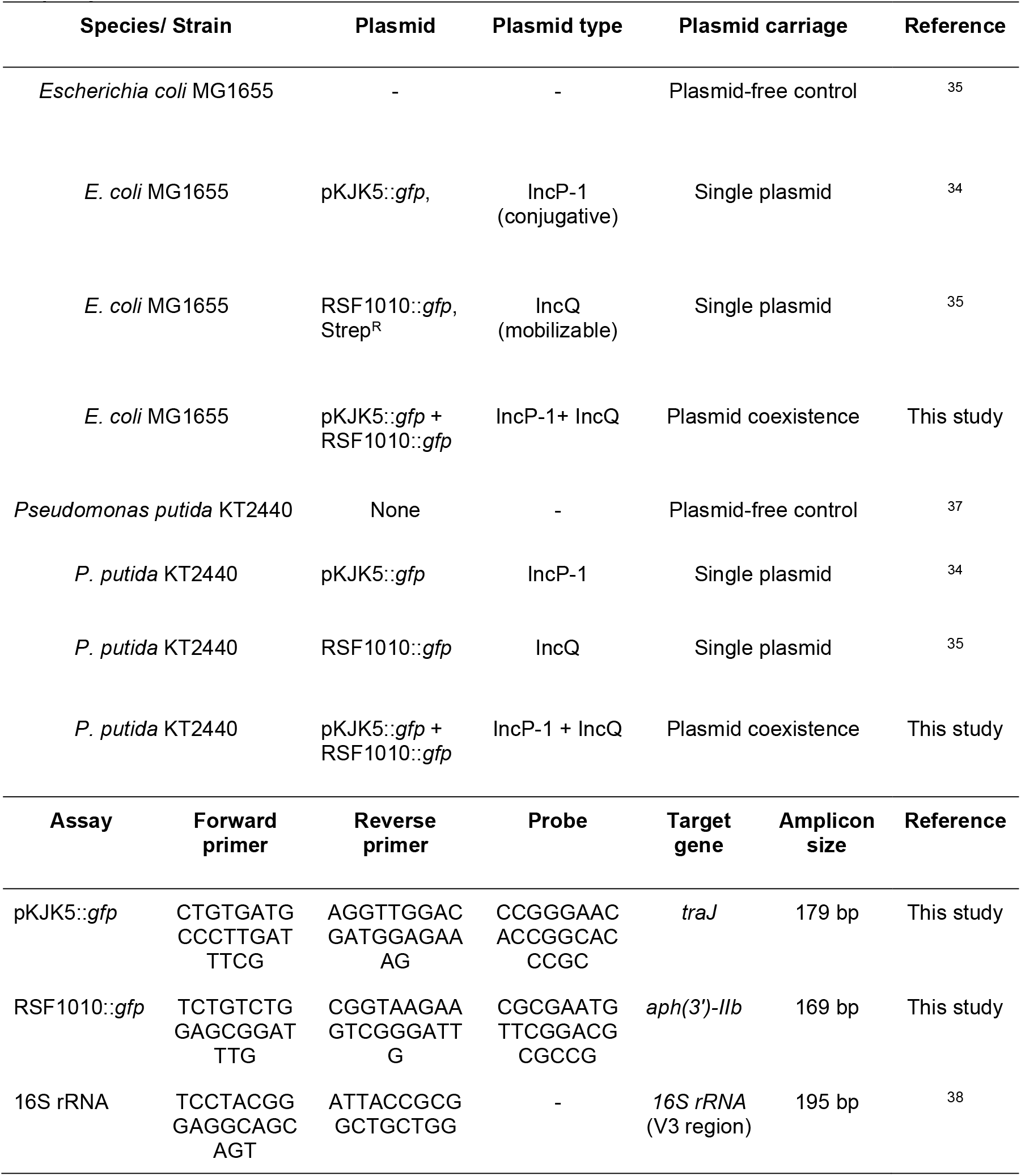
List of strains, plasmids, and primers used in this study. Km = Kanamycin, Tet = Tetracycline, Strep = Streptomycin.

### EV isolation from liquid cultures

EVs from each of the 8 strains (2 species x 4 plasmid configurations) were isolated from liquid bacterial cultures following a previously reported extraction protocol^39^ with minor modifications. Briefly, each strain was first grown overnight on LB agar plates containing antibiotics corresponding to plasmid-encoded resistance markers. A single colony was inoculated into 1 L of LB medium and incubated at 37°C with shaking at 120 rpm until the late exponential growth phase (OD_600_ = 1.0) was reached. Cultures were centrifuged at 10,000 × g for 30 minutes to remove bacterial cells, and the supernatant was sequentially filtered through 0.45 µm and 0.22 µm filters (GVS North America, USA). The filtrates were concentrated using a 100 kDa cutoff tangential flow filtration system (Vivaflow 200, Sartorius, Germany). To ensure complete removal of bacteria, the concentrates were additionally filtered through a 0.22 µm cellulose membrane filter (Merck Millipore, Cork, Ireland). EVs were pelleted by ultracentrifugation of the concentrate at 100,000 × g for 2 h at 4 °C using a SW32 Ti rotor (Beckman Coulter, USA), and resuspended in 200 μL PBS buffer. EV preparations were obtained from three independent biological replicates per condition (n=3). To ensure that DNA detected in subsequent analyses was exclusively associated with EVs, samples were treated with 2 U of Turbo DNase (Invitrogen, USA) for 30 minutes at 37°C. DNase was subsequently inactivated by heat treatment at 75°C for 15 minutes^40^. Genomic DNA and plasmid DNA extracted from *E. coli* carrying the pKJK5 plasmid served as positive controls to confirm the effectiveness of the DNase treatment. DNA concentrations were subsequently quantified using a Qubit 3.0 fluorometer with the dsDNA HS Assay Kit (Invitrogen, Thermo Fisher Scientific).

### Nanoparticle tracking analysis

Nanoparticle Tracking Analysis (NTA) was performed using a NanoSight LM10 instrument equipped with a 642 nm laser and a CCD camera (NanoSight Ltd., UK) to quantify the isolated EVs. Prior to sample analysis, instrument performance and focus were verified using 100 nm polystyrene latex standard beads (Malvern Panalytical). Samples were diluted in filtered 1× PBS (0.22 μm membrane) to adjust particle concentrations to the optimal measurement range of 1×10^7^–1×10^8^ particles/mL, and injected into the sample chamber with a sterile syringe. All measurements were conducted at a controlled temperature of 22°C. For each sample, a single 60-second video was captured. To minimize carryover between measurements, the sample chamber was flushed with ultrapure water prior to each injection. Data acquisition and processing were performed using NTA software version 2.0. All videos were analyzed using the following uniform settings: camera gain 1, detection threshold 12, blur size 3×3, minimum expected particle size 50 nm, and minimum track length 10. EV concentration (particles/mL) and size distribution were calculated from each video. These values were used to determine EV production per bacterial cell and plasmid copies per EV.

### Duplex Droplet Digital PCR of EV fractions for plasmid quantification

Duplex droplet digital polymerase chain reaction (ddPCR) was performed to simultaneously detect and quantify the two plasmid targets (pKJK5::*gfp* and RSF1010::*gfp*) in extracted EVs. ddPCR enables absolute quantification based on a most-probable-number approach and does not require external calibration standards^41^. ddPCR was carried out using the QX200 Droplet Digital PCR System (Bio-Rad, Hercules, CA, USA). Each 20 μL of reaction mix contained 10 μL of ddPCR Supermix for Probes (No dUTP, Bio-Rad), two primer pairs targeting a 179 bp conserved region of the pKJK5 plasmid and a 169 bp conserved region of the RSF1010 plasmid (final concentration of each primer 900 nM) and respective probes for pKJK5 labelled with a HEX fluorescence reporter and for RSF1010 labelled with a FAM reporter (final concentration 250 nM each) (Table 1). EV extracts were diluted 1:10 in nuclease-free water, and 5 μL of the diluted EV suspension was used as a template. QX200™ Droplet Generator and Droplet Generation Oil for Probes (both Bio-Rad) were used for droplet generation. The PCR amplification was performed in a 96-well plate (Bio-Rad, Hercules, CA, USA) on a C1000 Touch Thermocycler (Bio-Rad, Hercules, CA, USA), under the following cycling conditions: 95 °C for 10 minutes enzyme activation, 40 cycles of 94 °C for 30 seconds denaturation, 60 °C for 1 minute annealing, followed by a final hold at 98 °C for 10 minutes. The ramp rate was set at 2 °C/s. After amplification, the droplets were read using the QX200 Droplet Reader (Bio-Rad, Hercules, CA, USA). The data was analyzed using QuantaSoft software (Bio-Rad). This approach enabled absolute quantification of pKJK5 and RSF1010 copy numbers, which were subsequently used to calculate plasmid copies per EV. All ddPCR reactions were performed in technical duplicates for each independent biological replicate (n=3).

### Extraction of bacterial genomic DNA

To quantify intracellular plasmid copy numbers per bacterial cell, total genomic DNA was extracted from strains carrying either single plasmids or both coexisting plasmids. Total genomic DNA was extracted using the DNeasy PowerSoil Kit (Qiagen, Hilden, Germany) according to the manufacturer’s protocol, with modifications to sample preparation. Briefly, instead of 250 mg of solid sample, 2 mL of OD_600_=1 bacterial cultures grown under conditions identical to those used for EV extraction was centrifuged at 10,000 × g for 1 minute, and the resulting pellet was used for subsequent DNA extraction. DNA concentrations were quantified using a NanoDrop 2000 (Thermo Fisher, Waltham, MA, USA), and quality was assessed based on the 260/280 nm absorbance ratio, which ranged between 1.8 and 2.0 for all samples. Extracted DNA was stored at -20 °C until further analysis. Three independent biological replicates per condition were used for DNA extraction.

### Droplet Digital PCR with bacterial DNA

Plasmid copy number per bacterial cell was quantified using duplex ddPCR as described for EV extracts. DNA samples were normalized to 2 ng/μL and diluted 1:500 in nuclease-free water to avoid droplet saturation. A total of 5 μL of diluted DNA was used as a template. In addition, total cell numbers were quantified by ddPCR using the EvaGreen Assay targeting the 16S rRNA gene. Each 20 μL reaction contained 10 μL of ddPCR EvaGreen Supermix (Bio-Rad), the 331F/518R primer pair targeting a 195 bp region of the 16S rRNA gene (final concentration 250 nM) (Table 1)^38^, and 5 μL of the diluted template DNA. Droplet generation, thermal cycling, droplet reading, and data analysis were performed as described above. This approach enabled calculation of intracellular plasmid copy numbers per bacterial cell by normalizing plasmid abundance to total cell counts using the known content of seven 16S rRNA gene copies/cell in *E. coli* ^42^ and seven gene copies/cell in *P. putida*^43^.

### Data processing, visualization, and calculation of EV plasmid incorporation

Data analysis was performed in RStudio v. 2026.01.0+392^44^, and the R package ggplot2 v.3.5.1^45^ was used for data visualization. The analyzed variables included vesicle concentrations (vesicles per mL), vesicle production (vesicles per bacterial cell), plasmid copy numbers per vesicle, plasmid copy numbers per cell, and derived measures such as the fraction of intracellular plasmids incorporated into EVs and observed-to-expected plasmid ratios.

To estimate the fraction of the intracellular plasmid pool incorporated into EVs, total EV-associated plasmid copies were compared to total intracellular plasmid abundance in the cultures. Since EVs were isolated from 1 L of bacterial culture and resuspended in 200 μL, we first scaled all vesicle concentrations to the original culture volume by multiplying vesicle concentration in the EV suspension by the resuspension volume (0.2 mL). The total plasmid copies associated with EVs were then calculated by multiplying plasmid copy number per vesicle by the total number of vesicles in 1 L of the original culture. The intracellular plasmid abundance was estimated by multiplying plasmid copy number per cell by the number of cells per mL and scaling this value to 1 L.

The total intracellular plasmid abundance was calculated as the product of the plasmid copy number per cell and the cell density (cells mL^−1^) estimated from OD_600_ measurements. The fraction of plasmid in vesicles was thus calculated as:

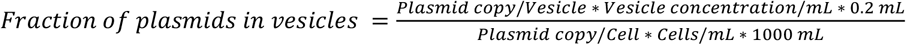

This fraction was then used to calculate expected plasmid copy numbers per vesicle under a stochastic packaging model, assuming proportional incorporation based on intracellular abundance:

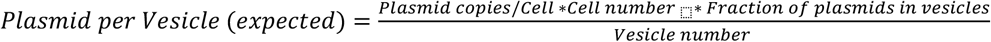

A linear model was fitted to log-transformed vesicle numbers to assess the effect of plasmid carriage on EV production. Plasmid carriage (no plasmid, single-plasmid, and dual-plasmid) was set as an explanatory variable. Similarly, a linear model was fitted to log-transformed intracellular plasmid copy numbers to assess the effects of host background, plasmid identity, and coexistence state, with these factors included as explanatory variables. The model included all main effects and their interactions. Effect sizes were estimated using partial eta squared (η^2^). A t-test was used to determine if plasmid incorporation in the vesicles with coexisting plasmids deviates from the stochastic model assuming proportional packaging based on single-plasmid EV packaging and intracellular plasmid abundance. Throughout, statistical significance was assumed for *P* < 0.05.

## Results

### Vesicle production is primarily determined by host background and not consistently influenced by plasmid carriage

We first assessed whether vesicle production differed between host backgrounds and plasmid carriage states. We thus quantified vesicle production of *E. coli* and *P. putida* strains carrying either: (i) no plasmids, (ii) pKJK5 or (iii) RSF1010 individually, or (iv) coexisting RSF1010 and pKJK5 plasmids. Using NTA, we detected vesiculation in all tested strains (Fig.1A).

To quantify the contribution of host and plasmid effects, we fitted a linear model on log-transformed vesicle production data, including host background, pKJK5 presence, and RSF1010 presence as factors. This analysis identified host background as the dominant driver of vesicle production (η^2^ = 0.42, P < 0.0001), with *E. coli* producing significantly more vesicles per cell (1.6 - 4.8 × 10^-3^) than *P. putida* (0.3 - 2.6 × 10^-3^). Plasmid carriage-related effects were variable and strongly host-dependent (Fig. 1A). While pKJK5 showed a moderate effect in *E. coli*, RSF1010 was associated with increased vesicle production in *P. putida*, as reflected by significant host– plasmid interaction terms (η^2^ = 0.25, P < 0.0001). However, these effects were not consistent across hosts and did not represent a generalizable influence of plasmid carriage on vesicle production.

**Figure 1.**
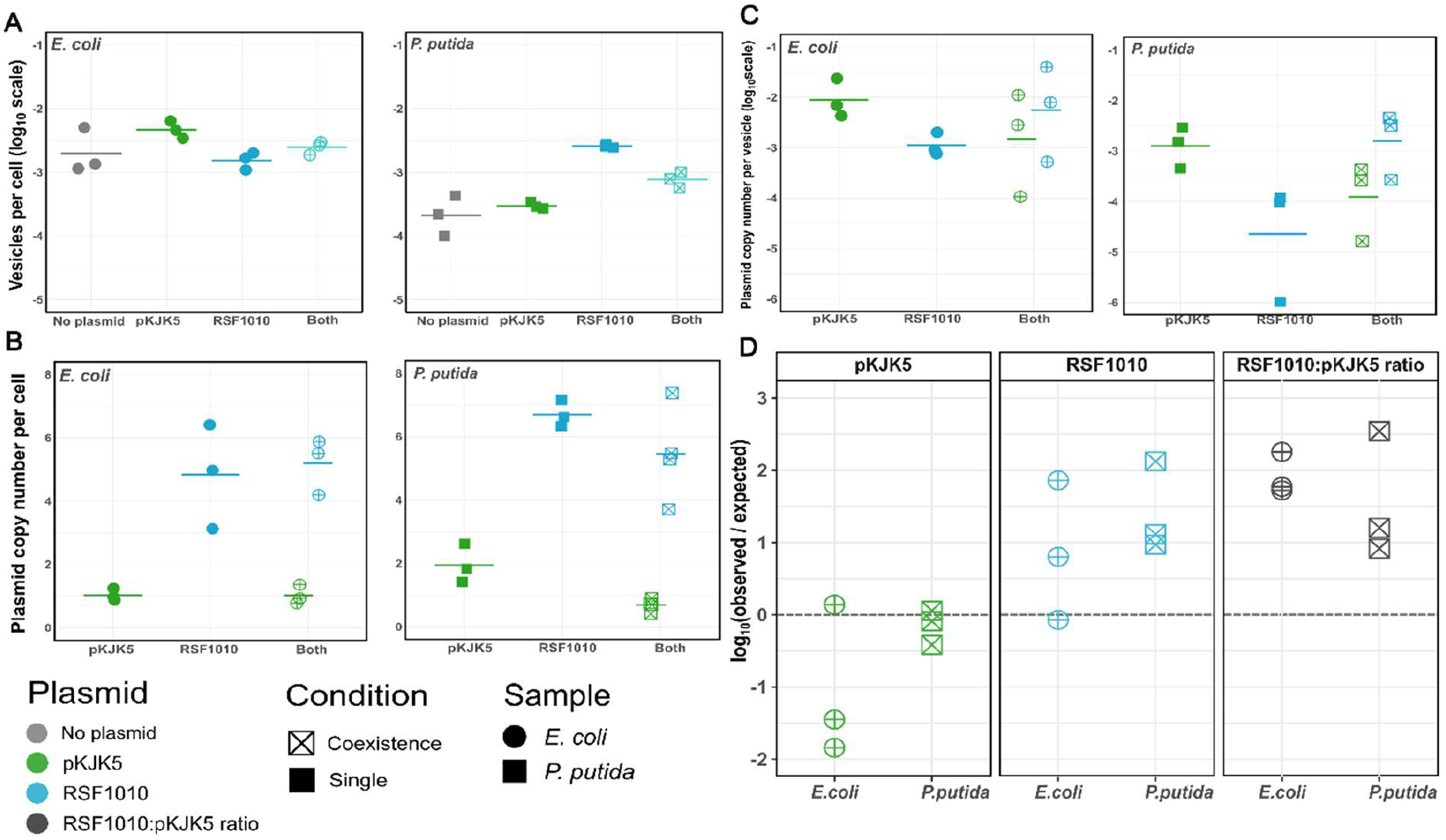
Effects of plasmid coexistence on vesicle production and plasmid incorporation into EVs produced by *E. coli* and *P. putida*. (A) Log_10_-transformed number of EVs produced per cell by *E. coli* and *P. putida* strains with no plasmid, a single plasmid, and dual plasmid carriage. (B) Intracellular plasmid copy number per cell in strains harboring single or coexisting plasmids. (C) Log_10_-transformed plasmid copy number per EV. (D) Log_10_ -transformed ratio of observed to expected EV-associated plasmid abundance. The dashed line indicates the stochastic expectation of plasmid incorporation into EVs (log_10_(observed/expected) = 0). Expected values were derived from plasmid incorporation observed under single plasmid conditions and intracellular plasmid abundance observed under plasmid coexistence.

Together, these results indicate that vesicle production is primarily governed by host physiology, while plasmid effects are context-dependent and comparatively minor under the tested conditions. This indicates that differences in vesicle production alone cannot explain differences in EV plasmid-cargo composition observed below.

### Intracellular plasmid abundance is stable and defines expected packaging ratios

To establish the baseline intracellular plasmid abundance, we quantified plasmid copy number per cell for both plasmids in single-plasmid and coexisting conditions across both bacterial hosts. As expected, the high-copy-number plasmid RSF1010 was consistently more abundant than the low-copy-number plasmid pKJK5 across all conditions (Fig. 1B).

A linear model on log-transformed copy numbers, including host background, plasmid identity, and coexistence state, identified plasmid identity as the dominant determinant of intracellular abundance (η^2^ = 0.84, partial η^2^ = 0.93, P < 0.0001). In contrast, host background had a significant but small effect (η^2^ = 0.003, partial η^2^ = 0.05, P < 0.05). Coexistence alone did not significantly affect intracellular copy number overall (η^2^ = 0.03, partial η^2^ = 0.34, P = 0.9) but a significant interaction between host and coexistence was observed (η^2^ = 0.03, partial η^2^ = 0.36, P < 0.01), indicating coexistence effect on plasmid copy number was host-dependent. No significant other interaction terms were detected (all P > 0.05).

Consistent with this, in *E. coli*, pKJK5 copy number remained essentially unchanged between single-plasmid and coexisting conditions (1.03 ± 0.19 vs 1.02 ± 0.30 copies per cell), while RSF1010 showed only a minor increase (4.83 ± 1.65 vs 5.19 ± 0.89). In *P. putida*, pKJK5 decreased from 1.95 ± 0.61 to 0.69 ± 0.21 copies per cell under coexistence, whereas RSF1010 decreased more moderately from 6.70 ± 0.42 to 5.46 ± 1.50. Accordingly, the intracellular RSF1010:pKJK5 ratio was approximately 5:1 in *E. coli* under both single and coexisting conditions, and increased from approximately 3.8:1 to 7.9:1 in *P. putida* under coexistence.

Together, these results show that intracellular plasmid abundance is primarily defined by plasmid identity and does not show consistent shifts across hosts. The intracellular ratios of the two plasmids, therefore, provide the null expectation for stochastic EV packaging tested below.

### Conjugative plasmids are preferentially incorporated into EVs in isolation

We next quantified plasmid incorporation into EVs under single plasmid conditions (Fig. 1C). Across both hosts, the conjugative plasmid pKJK5 was consistently enriched in EVs relative to the mobilizable plasmid RSF1010, despite its lower intracellular abundance and larger size.

In *E. coli*, pKJK5 reached 1.2 ± 1 × 10^-2^ copies per vesicle, approximately 10-fold higher than RSF1010 (1.2 ± 0.7 × 10^-3^). In *P. putida*, this enrichment was even more pronounced, with pKJK5 exceeding RSF1010 by approximately 23-fold. Given the higher intracellular abundance of RSF1010, stochastic packaging would predict its dominance in EVs, with expected RSF1010:pKJK5 ratios of approximately 5:1 in *E. coli* and 3.8:1 in *P. putida*. Instead, the observed ratios were reversed, with pKJK5 significantly dominating EV cargo above stochastic expectation in both hosts (t-test, P < 0.001).

Thus, under single-plasmid conditions, EV-associated plasmid composition does not reflect intracellular abundance but instead shows a consistent enrichment of the conjugative plasmid across host backgrounds. These findings support the hypothesis that plasmid-specific properties, such as membrane association potentially linked to conjugative transfer machinery, promote preferential incorporation of conjugative plasmids into EVs.

However, natural bacterial systems frequently involve co-resident plasmids, where interactions between plasmids can alter access to membrane-associated transfer machinery. We therefore next tested whether such interactions reshape plasmid incorporation into EVs beyond patterns observed under single-plasmid conditions.

### Plasmid co-residence reverses EV cargo composition and deviates from stochastic expectations

To test whether plasmid–plasmid interactions alter EV cargo composition, we quantified plasmid incorporation into EVs under coexisting conditions and compared observed values to expectations based on intracellular plasmid abundance. Under plasmid co-residence, EV cargo composition shifted markedly in both hosts. In contrast to single-plasmid conditions, the mobilizable plasmid RSF1010 became dominant in EVs, exceeding pKJK5 by approximately 4-fold in *E. coli* and 11-fold in *P. putida* (Fig. 1C).

In absolute terms, RSF1010 increased from 1.2 ± 0.7 × 10^-3^ to 1.6 ± 1 × 10^-2^ copies per vesicle in *E. coli*, corresponding to an approximately 13-fold enrichment under coexistence. In *P. putida*, RSF1010 increased from 7.3 ± 6 × 10^-5^ to 2.7 ± 2 × 10^-3^ copies per vesicle, corresponding to an approximately 39-fold enrichment. In contrast, pKJK5 decreased from 1.2 ± 1 × 10^-2^ to 5 ± 5 × 10^- 3^ copies per vesicle in *E. coli* (approximately 2.4-fold reduction), and from 1.7 ± 1.2 × 10^-3^ to 2.4 ± 2 × 10^-4^ in *P. putida* (approximately 7-fold reduction).

Species-specific analyses showed consistent directional shifts in EV-associated plasmid ratios. Based on the intracellular plasmid abundance and single-plasmid EV packaging patterns above, expected RSF1010:pKJK5 ratios would have remained approximately 0.10 in *E. coli* and 0.04 in *P. putida*. Instead, under coexistence, the observed ratios increased to approximately 4 and 11, respectively.

Comparison to stochastic expectations revealed that RSF1010 was significantly enriched in EVs relative to predicted values across both species (one-sample t-test, P < 0.01), whereas pKJK5 showed a consistent reduction (P = 0.14). Consequently, the EV-associated RSF1010:pKJK5 ratio significantly deviated from the expected ratio (P < 0.001) (Fig. 1D). Under a stochastic packaging model, EV-associated plasmid composition would be expected to reflect intracellular abundance and the packaging bias observed under single-plasmid conditions. Instead, plasmid co-residence shifted EV cargo toward dominance of the mobilizable plasmid.

Together, these results demonstrate that plasmid incorporation into EVs is non-random and is actively reshaped by plasmid–plasmid interactions, supporting the second hypothesis. This interaction-driven restructuring of EV cargo was conserved across both hosts, despite strong differences in vesicle production and host-specific plasmid effects. This decoupling demonstrates that EV cargo composition is not governed by vesicle production dynamics, but instead by selective processes linked to plasmid-specific properties and their interactions. Thus, plasmid– plasmid interactions define EV-mediated gene transfer outcomes independently of vesicle abundance.

## Discussion

In this study, we investigated whether principles governing plasmid transfer, including plasmid mobility type and plasmid–plasmid interactions^13,25,32^, extend beyond conjugation to influence their incorporation into EVs. Specifically, we tested whether conjugative and mobilizable plasmids differ in their propensity for EV incorporation, and whether plasmid co-residence alters this pattern through interactions between coexisting plasmids. Whereas both plasmids were detected in EVs, the EV fraction with pKJK5 as cargo dominated over that with RSF1010 in strains carrying individual plasmids. In contrast, under coexistence, this relationship was reversed, with RSF1010 enriched and pKJK5 depleted in EV cargo. These effects were consistent across both bacterial hosts. Together, these findings demonstrate that plasmid incorporation into EVs is not solely determined by intracellular abundance but is shaped by plasmid-specific properties and their interactions, extending plasmid transfer dynamics beyond classical conjugation.

The preferential incorporation of pKJK5 under single-plasmid conditions diverges from its lower intracellular abundance relative to RSF1010, challenging the assumption that EV packaging is driven by plasmid copy number alone^10,27^. A plausible explanation is the spatial localization of plasmids within the bacterial cell. Similar to other conjugative plasmids, pKJK5 encodes a type IV secretion system (T4SS) that enables its autonomous transfer between bacteria^13,28^. Autonomous transfer requires assembly of the T4SS, including coupling proteins and associated components that localize the plasmid-relaxosome complex at the cell membrane^46,47^, thereby positioning plasmid DNA in proximity to the cell envelope. In contrast, mobilizable plasmids such as RSF1010 lack the MPF^34,48^. Although mobilizable plasmids can associate with the membrane, for example via MobB protein of the RSF1010 plasmid^49^, their association with the cell membrane is likely less structured than that of conjugative plasmids. As EVs originate from the bacterial membrane, the incorporation of the components that are more consistently localized at the membrane is expected to be more efficient, which may favour conjugative over mobilizable plasmids.

In contrast, mobilizable plasmids such as RSF1010 lack the MPF system34,48 and therefore do not encode a dedicated transfer apparatus for organized membrane localization. Although mobilizable plasmids can associate with the membrane, for example via the MobB protein of RSF1010, this association is likely less structured than that of conjugative plasmids. As EVs originate from the bacterial membrane, they are expected to preferentially capture components that are membrane-proximal yet dynamically accessible, which may favour mobilizable over conjugative plasmids.

An additional, but likely secondary, mechanism could involve local bacterial outer membrane disturbance^50^ associated with T4SS assembly and pilus formation^51^, which has been proposed to promote vesiculation. However, in our data, plasmid carriage did not lead to consistent increases in EV production across hosts, indicating that plasmid-induced membrane perturbation is not a dominant driver of EV formation. Given that EV production is largely governed by host background and likely includes contributions from both membrane blebbing or cell lysis^4–6^, the observed enrichment of pKJK5 occurs despite this heterogeneous EV background. Importantly, the proposed mechanism of membrane proximity does not depend on a specific mode of EV biogenesis, as vesicles derived from either membrane blebbing or cell lysis are equally expected to selectively capture membrane-associated material^5^. Together, these observations support a model in which spatial proximity to membrane-associated transfer machinery, rather than EV production per se, drives preferential incorporation of conjugative plasmids into EVs.

Plasmid co-residence can influence access to membrane-associated transfer machinery, as mobilizable plasmids exploit conjugative systems and coexisting plasmids can interfere with each other’s transfer^25^. In contrast to single-plasmid conditions, coexistence reversed EV cargo composition, suggesting that interactions between conjugative and mobilizable plasmids reshape access to sites of preferential EV cargo incorporation. In multi-plasmid conjugative systems, mobilizable plasmids are known to exploit the transfer machinery encoded by conjugative plasmids, and RSF1010 is a well-characterized example that is efficiently mobilized by IncP-1 plasmids such as pKJK5^34,36,52^. This process relies on the affinity between the relaxosome of the mobilizable plasmid and the type IV coupling protein (T4CP) complex of the conjugative system, which determines access of RSF1010 to membrane-associated transfer sites and thereby its mobilization efficiency^53^. Under coexistence, such interactions may lead to preferential recruitment of RSF1010 to these membrane-associated sites, increasing its proximity to the cell envelope and thereby its likelihood of incorporation into EVs. At the same time, this recruitment is expected to reduce the availability of T4CPs and associated transfer complexes for pKJK5 itself, resulting in its relative depletion from membrane-associated regions and consequently from EV cargo, as has been shown for other conjugative-mobilizable plasmid pairs^54,55^. Thus, competition for and preferential access to membrane-associated transfer machinery can simultaneously explain the enrichment of RSF1010 and the depletion of pKJK5 observed under plasmid coexistence. These findings indicate that interactions between mobilizable and conjugative plasmids govern not only direct gene transfer through conjugation but also which genetic elements enter the extracellular gene pool through EVs.

Despite pronounced differences in EV production observed between hosts, the direction of plasmid packaging dynamics was conserved. This indicates that host background primarily modulates EV abundance and effect size, while plasmid–plasmid interactions determine cargo composition. Host physiology may nonetheless influence the ecological relevance of this mechanism. In *P. putida*, EV production is more strongly associated with stress responses and membrane instability, whereas in *E. coli*, vesiculation occurs at a higher basal level^56–58^. Under stress conditions, which represent windows of bacterial adaptation where plasmid-encoded traits can provide strong selective advantages^17,59,60^, increases in EV production may amplify the interaction-driven restructuring of plasmid cargo observed here, thereby contributing more substantially to extracellular gene flux. In complex natural microbial communities, where both stress exposure and multi-plasmid carriage are common, these dynamics may influence not only which plasmids are transferred, but also which host populations contribute to EV-mediated gene pools. Given that mobilizable plasmids are often overrepresented in natural microbial communities^25,61–63^, which can be related to their smaller size, lower intrinsic costs, and ability to utilize conjugative plasmids for transfer^60,64,65^, the plasmid–plasmid interactions described here are likely widespread. Our results suggest that such interactions extend beyond conjugation to shape the composition of extracellular genetic material, with potential consequences for the dissemination of AMR genes and other adaptive traits.

Despite these insights, this study has several limitations. First, we focused on a single pair of model plasmids, and different combinations of conjugative and mobilizable plasmids may exhibit distinct interaction dynamics and packaging behavior. Second, the experiments were conducted in two bacterial hosts under controlled laboratory conditions, and vesicle-mediated packaging patterns may differ in more complex microbial communities. Third, the proposed mechanism linking plasmid interactions to membrane-associated positioning and EV incorporation remains inferred rather than directly demonstrated. Future work should therefore expand to additional plasmid combinations and incompatibility groups, including different plasmid variants and targeted mutants with altered mobilization or coupling functions ^66^, to directly test the role of conjugation machinery in mediating plasmid recruitment into vesicles. Direct visualization of plasmid localization relative to the cell envelope, for example, using fluorescence-based approaches^67^, would provide critical evidence for spatial positioning as the suggested driver of EV incorporation. Beyond controlled systems, extending these analyses to more complex microbial communities will be essential to determine whether the interaction-driven restructuring of EV cargo observed here persists under environmentally relevant conditions. Finally, assessing the functional consequences of EV-mediated plasmid transfer, including uptake and establishment in recipient bacteria and their rates relative to established HGT mechanisms, will be critical to evaluate its ecological and evolutionary significance.

In summary, our results demonstrate that plasmid incorporation into extracellular vesicles is not governed by stochastic processes or intracellular abundance alone, but is shaped by plasmid-specific properties and their interactions. Conjugative plasmids are preferentially incorporated under single-plasmid conditions, while plasmid co-residence reverses this pattern through interaction-driven restructuring of access to membrane-associated transfer machinery. These findings extend the role of plasmid–plasmid interactions beyond conjugation and identify them as a key determinant of extracellular gene transfer. Given the ubiquity of vesiculation and multi-plasmid carriage in microbial communities, this mechanism represents a previously unrecognized route by which plasmid interactions shape the dissemination of antibiotic resistance and other adaptive traits.

## Author Contributions - CRediT

**Faina Tskhay:** Conceptualization; Methodology; Validation; Formal analysis; Investigation; Data Curation; Writing - Original Draft; Writing - Review & Editing; Visualization

**Haining Huang:** Conceptualization; Methodology; Formal analysis; Investigation; Writing - Review & Editing; Visualization; Funding acquisition

**Robin Starke:** Methodology; Formal analysis; Data Curation; Writing - Review & Editing

**Magali de la Cruz Barron:** Methodology; Writing - Review & Editing

**Maria Pilar Garcillán Barcia:** Writing - Review & Editing

**Thomas U. Berendonk:** Conceptualization; Resources; Writing - Review & Editing; Visualization; Supervision; Project administration; Funding acquisition

**Anja Worrich:** Conceptualization; Resources; Writing - Review & Editing; Supervision; Project administration; Funding acquisition

**Uli Klümper:** Conceptualization; Methodology; Resources; Data Curation; Writing - Original Draft; Writing - Review & Editing; Visualization; Supervision; Project administration; Funding acquisition

## Acknowledgments

The authors thank Christiane Zschornack, Melanie Tannert, and Steffen Kunze for technical support in the laboratory.

## Data Availability

The datasets supporting the conclusions of this article are included within the article and its additional files or available through the corresponding author upon reasonable request.

